# Bayesian Hypothesis Testing And Experimental Design For Two-Photon Imaging Data

**DOI:** 10.1101/410357

**Authors:** Luke E. Rogerson, Zhijian Zhao, Katrin Franke, Philipp Berens, Thomas Euler

## Abstract

Variability, stochastic or otherwise, is a central feature of neural circuits. Yet the means by which variation and uncertainty are derived from noisy observations of neural activity is often unprincipled, with too much weight placed on numerical convenience at the cost of statistical rigour. For two-photon imaging data, composed of fundamentally probabilistic streams of photon detections, the problem is particularly acute. Here, we present a complete statistical pipeline for the inference and analysis of neural activity using Gaussian Process Regression, applied to two-photon recordings of light-driven activity in *ex vivo* mouse retina. We demonstrate the flexibility and extensibility of these models, considering cases with non-stationary statistics, driven by complex parametric stimuli, in signal discrimination, hierarchical clustering and inference tasks. Sparse approximation methods allow these models to be fitted rapidly, permitting them to actively guiding the design of light stimulation in the midst of ongoing two-photon experiments.

## Introduction

Over the last two decades, two-photon (2P) imaging has become one of the premier tools for studying coding and computations in neural systems from the population level down to individual neural compartments [1]. The resulting data is highly variable due to the inherent variability of neurons and technical sources of noise in the imaging process [2, 3]. Yet we typically assume that there is a smooth latent function describing the activity of a neuron or a neural compartment. In a typical analysis pipeline for 2P data, we attempt to recover this function by grouping noisy observations from pixels into regions of interest (ROIs), which covers the soma or a different compartment of a neuron, temporally interpolating them to a common frame rate and averaging across repetitions of the same stimulus. Each stage is intended to smooth the observations and get closer to the “true” underlying activity function of the neuron. To measure the uncertainty about this latent activity function, often the inter-repetition variability is used, with little assessment of whether this reflects the signal variability prior to the grouping and smoothing operations.

Here, we propose a different approach based on Gaussian Process (GP) regression [4] to infer signals from 2P recordings in a statistically principled manner, propagating the uncertainty all the way from the measurements to the desired inference about the latent activity function. Gaussian processes are a family of Bayesian models which form probabilistic representations of functional signals. In contrast to typical pre-processing pipelines (Fig. 1a), the statistical properties of the observation set are considered explicitly as part of the model optimisation (Fig. 1b). Recently developed sparse GP approximations allow us to apply these models to comparatively large datasets with several thousand observations, as are common in 2P experiments.

**Fig. 1:**
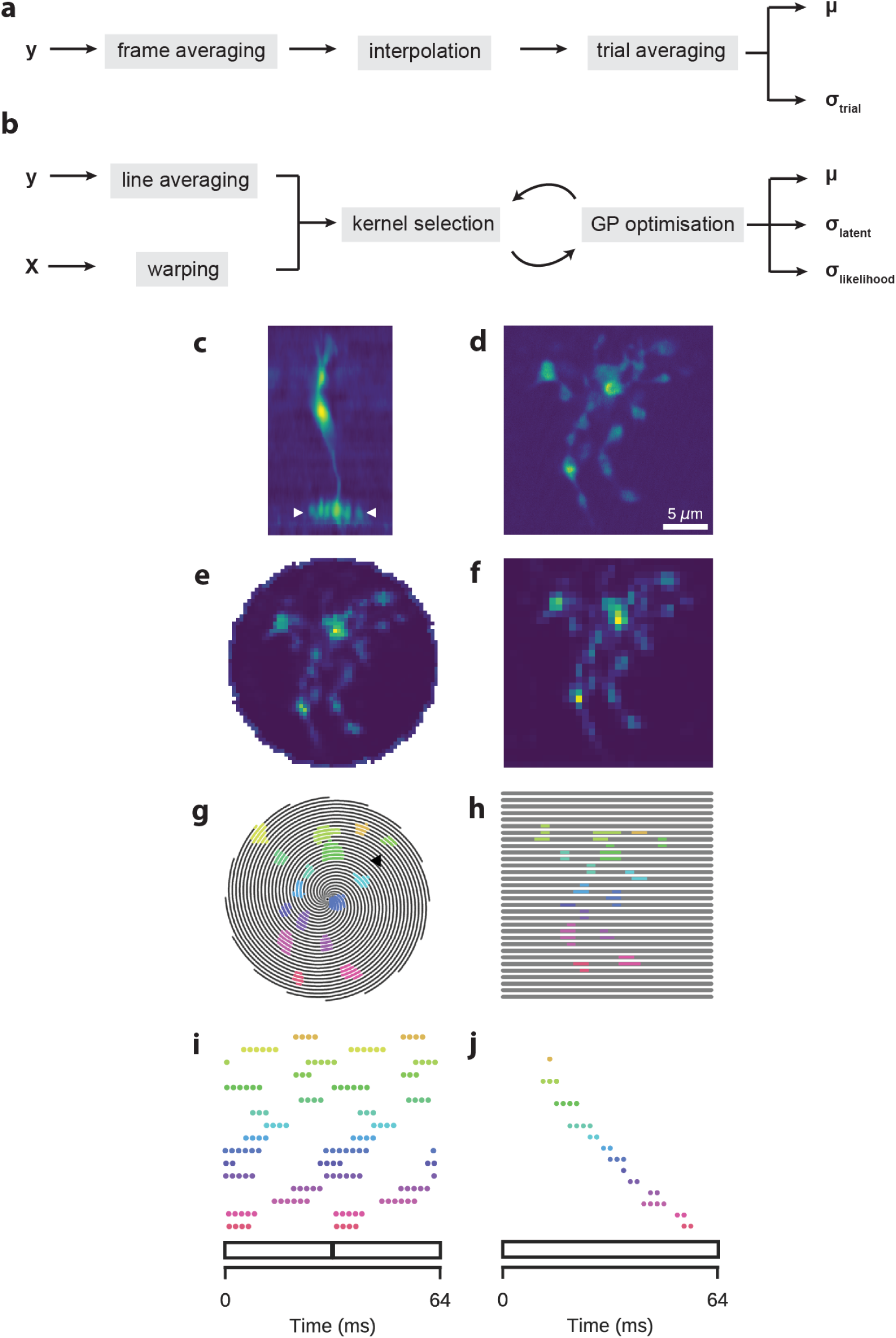
Two-Photon Imaging Of Retinal Neurons. Data: Retinal bipolar cell filled with OGB-1 via sharp electrode injection and recorded using linear and spiral scan configurations. (**a**) Typical pre-processing pipeline. Two-photon imaging data y is averaged over a recording frame, interpolated to a high temporal resolution to compute a mean signal μ and an inter-trial variance σ_trial_. (**b**) Proposed pre-processing pipeline using GP regression. Two-photon imaging data is averaged over scan lines within a recording frame, a kernel is selected and the GP is optimised using both the imaging data and the stimulus features X. Stimulus features may be warped to allow for non-stationary autocorrelation. Kernel selection and optimisation may be an interactive process. The resulting model consists of a mean signal μ, and the variance with (σ_likelihood_) and without (σ_latent_) likelihood noise. (**c**) Vertical profile. Image coloured according to fluorescence intensity (yellow: high, blue: low). (**d**) Horizontal (x-y), high-resolution scan of axon terminal system (512 x 512 pixels), corresponding to the domain between the two white ticks in (c). (**e**) Spiral scan of axon terminal system (16 spirals; 31.25Hz), as above. (**f**) Linear scan of axon terminal system (32 lines; 15.625Hz), as above. (**g**) Spiral scan trajectory with ROI mask superimposed. Black lines indicate scan trajectory. Colours correspond to discrete ROIs. (**h**) As (e), but with a linear scan configuration. The same ROIs were used. (**i**) Time points at which the scan trajectory intersects with the ROI mask in (e). 64ms span corresponds to two spiral scan frames. (**j**) As (**g**), but for a linear scan configuration. ROIs correspond to those in (f). 64ms span corresponds to one linear scan frame.

Using 2P recordings of calcium and glutamate dynamics in isolated mouse retina, we show how these models can be used to construct probabilistic representations of neural activity. We treat several use cases, in which we also demonstrate the superiority of our framework over standard approaches: First, we show that GP-based analysis of 2P recordings can be used to perform principled comparisons between the responses of a given cell under different conditions, allowing one to identify parts of the response where cells responses change significantly. Second, we show that this can be exploited to perform a hierarchical clustering of cell responses and provides quantitative criteria for deciding how many clusters to keep. In addition, we show how the framework can be used to test which stimulus parameters influence neural activity in an ANOVA-like framework. Finally, we show how the representation of uncertainty can be exploited for experimental design, informing the choice of parameters to optimally reduce the uncertainty about the neural response.

## Results

We applied a Bayesian framework based on Gaussian Processes to efficiently infer latent neural activity with uncertainty estimates from recordings of light stimulus-driven activity in the mouse retina. The retina decomposes a stream of images into a set of parallel channels representing salient stimulus features. The central circuit of this network is a feedforward pathway relaying the initial signal from the photoreceptors through the intermediate bipolar cells to retinal ganglion cells (RGCs), and from there through the optic nerve to the rest of the visual system. Inhibitory interneurons called horizontal and amacrine cells play key roles in the adaptation and feature extraction (for review, see [5]). In the datasets analysed here, we measured three stages of the excitatory pathway: Firstly, the presynaptic calcium signal in the axon terminals of a bipolar cell using the synthetic indicator dyes Oregon-Green BAPTA-1 (OGB-1) and GCaMP6f (the latter data previously published in [6]. Secondly, the glutamate release from these terminals, as measured by the genetically-encoded biosensor iGluSnFR [7, 8]. Finally, the calcium signal in RGC somata loaded with OGB-1 through bulk electroporation [9].

In our framework, a GP model infers a latent activity function from the observed signal. In addition, it estimates the associated uncertainty including likelihood or observation noise, and the latent uncertainty once the likelihood noise is removed. One can intuitively understand the latter as the uncertainty about the activity function given the observations – the quantity we care about when performing statistical inference. While this would reduce to zero in the limit of infinite observations, the likelihood noise component will persist, as it derives from a combination of biological, measurement and numerical noise sources which do not depend on the size of the observation set.

### Modelling Uncertainty Using Gaussian Processes

Our first objective was to infer the latent activity function and its associated uncertainty from our observations of the activity of single ROIs, located on individual synaptic axon terminals of a bipolar cell. We applied the GP models to bipolar cell calcium and glutamate signals measured in such ROIs during the presentation of a spatially homogeneous light stimulus including a light step and variations in temporal frequency and contrast (chirp stimulus), as used in previous studies [6, 10] (Fig. 2a).

**Fig. 2:**
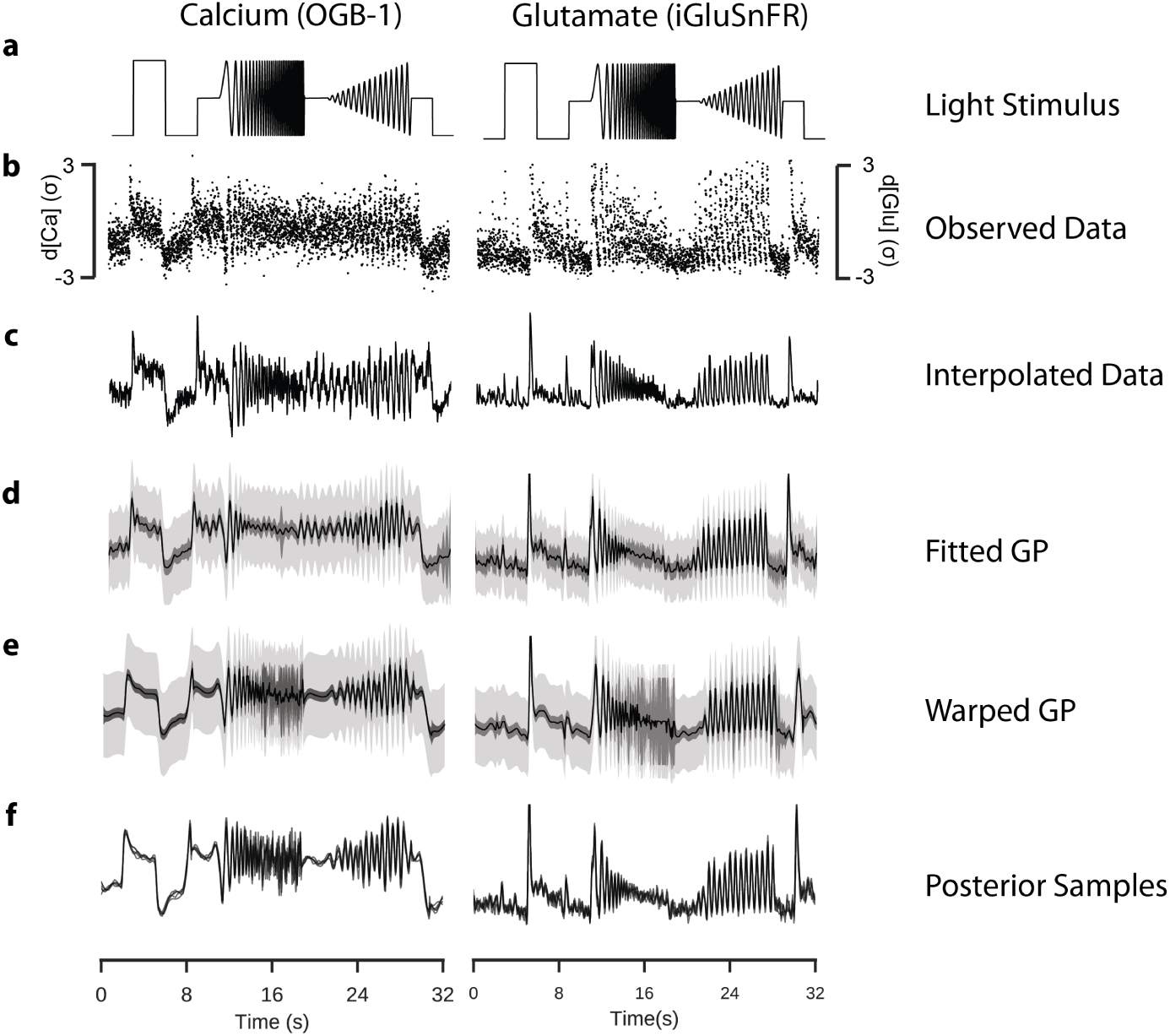
Inference Of Signals From Two-Photon Data. Data: ROI from a retinal bipolar cell filled with OGB-1 via sharp electrode injection (left), and a different ROI from a scan field with bipolar cell terminals in a retina expressing iGluSnFR (right); both recorded using spiral scan configurations. Model: RBF kernel, 300 inducing inputs, 25 iterations per fit, best of 6 fits per model. (**a**) “Chirp” light stimulus, consisting of a light step, a frequency-modulated sine wave and a contrast-modulated sine wave. Full field. (**b**) Observed activity of a single ROI. Each point corresponds to the mean activity of the ROI in a single scan line. (**c**) Estimate of underlying signal from frame averaging, cubic spline interpolation and averaging over trials. This corresponds to the typical approach used in previous papers [6]. (**d**) Fitted sparse Gaussian process. Black line indicates the mean signal. Intervals indicate uncertainty of the signal with and without the additive likelihood noise (light and dark grey, respectively), to 3 standard deviations. (**e**) Fitted sparse warped Gaussian process. Input warping uses the warping function shown in the following figure. Model has been projected back onto the original time axis. (**f**) Five posterior samples drawn from the fitted sparse warped Gaussian process models.

In bipolar cells injected with the calcium indicator OGB-1, it is possible to image the anatomy of the cell by recording 3D stacks of x-y images at regular intervals along the vertical (z) axis (Fig. 1c). High resolution scans allowed us to identify individual axon terminals (Fig. 1d). Faster scans with lower spatial resolution are required to resolve neural activity, although the required reduction in resolution is substantially less for spiral configurations relative to classical linear configurations (Fig. 1e,f). From recordings of two different pieces of retinal tissue, one labelled with OGB-1 and the other with iGluSnFR, we used the observed activity of a ROI (Fig. 2b), and inferred a signal for each repeat using frame-averaging and cubic-spline interpolation (Fig. 2c), corresponding to the classical way of inferring these functions (i.e. [6, 10]).

We then fitted a GP with a radial basis function (RBF) kernel on the time axis to the observed activity (Fig. 2d). We monitored the computation time and calculated the likelihood of an out-of-sample test set to determine a suitable number of inducing inputs. Surprisingly, this indicated that there was already little improvement in the performance of the model when more than 250 data points were used (Supplementary Fig. S1a,b), and that relatively few iterations of the fitting algorithm were required (Supplementary Fig. S1c).

To account for temporal non-stationarities in the neural response, we then compared the sparse GP model to an extended model with input warping. One assumption of classical GP models is that the function space has a stationary autocorrelation function, i.e. that its correlational structure does not change with respect to a predictor, such as time. However, light induced neural activity like responses to the chirp, which have highly non-stationary correlational structure, are likely to show a commensurate non-stationarity in the response. We computed a warping function which transforms the time dimension such that the stimulus input has a stationary autocorrelation structure. We then used this warping function to transform the input to the GP model of the response, under the assumption that the correlational structure of the response matched that of the stimulus input [11, 12](Fig. 3b). By performing this extra processing step, we were able to fit a model which could vary in its autocorrelation.

**Fig. 3:**
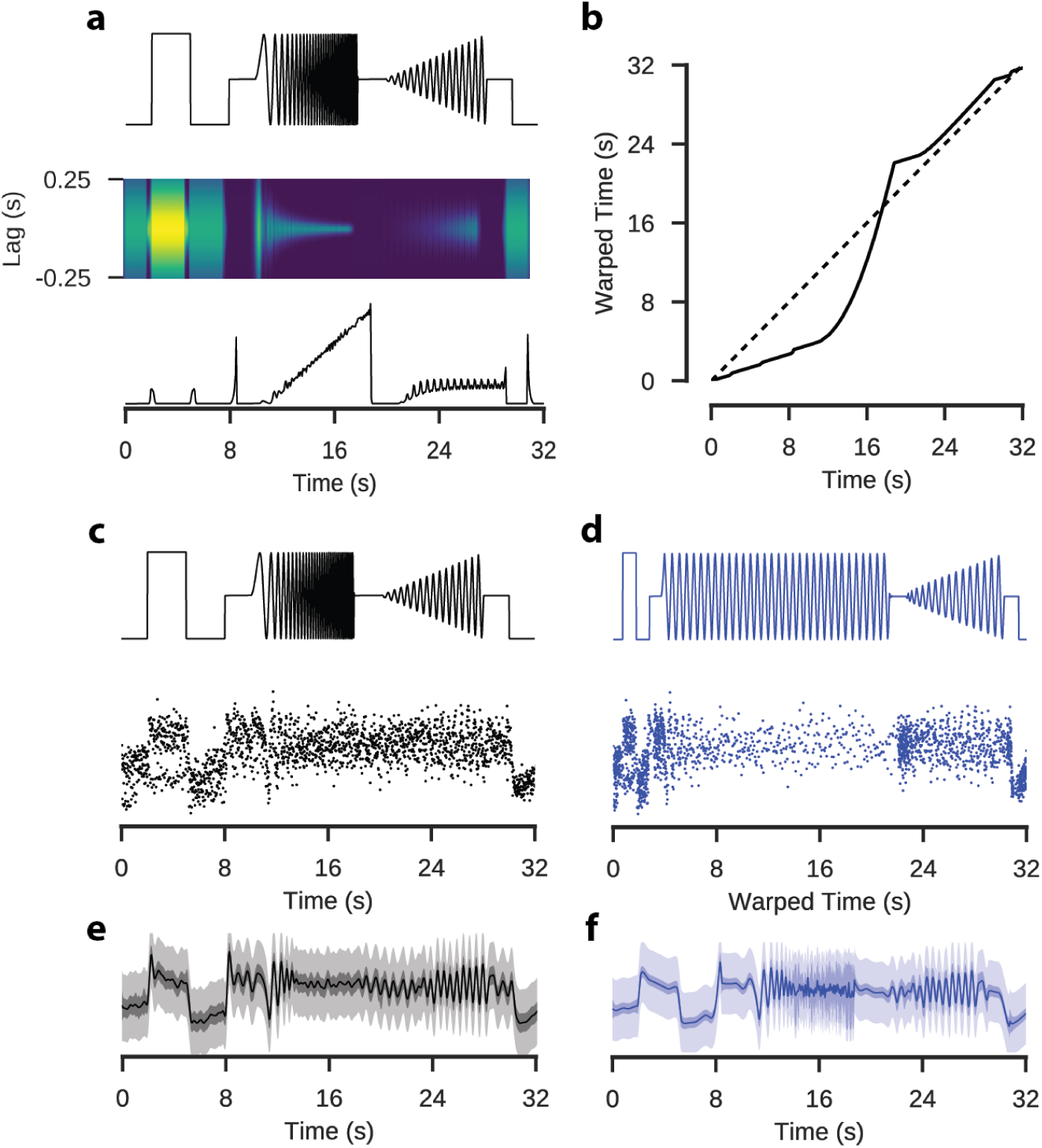
Application Of A Warping Function To Model Input Features. Data: ROI from a retinal bipolar cell filled with OGB-1 via sharp electrode injection. Model: RBF kernel, 300 inducing inputs, 20 iterations per fit, best of 3 fits per model. (**a**) “Chirp” stimulus (top). Auto-correlation functions corresponding to Gaussian curves fit to the empirical autocorrelation function over a 500ms window (middle). Lengthscale parameter of the Gaussian distribution fitted to the autocorrelation functions. (**b**) Cumulative distribution of the inverse lengthscale over time. If the signal were stationary, the lengthscale would be constant, corresponding to the dashed line. This cumulative distribution maps time onto a warped time dimension. (**c**) Full field Chirp stimulus with observations of the activity of one ROI labelled with OGB-1. (**d**) The same stimulus and observations after a warping operation has been applied. (**e**) GP fitted to the original data. (**f**) GP fitted to the warped data. The function has been projected back onto un-warped time. Note the increased uncertainty in regions where the stimulus is changing rapidly and the variations in the smoothness of the inferred signal over time.

Our results show a clear difference between the predictions of the warped GP model and the simpler stationary one. In the simpler model, the selected parameters reflect a trade-off between models which fit closely to each of the different stimulus components (i.e. steps vs. chirps), resulting in an inferred mean signal which appears noisy during the light step and poorly tracks the faster chirp oscillations. As a consequence, the inferred uncertainty was relatively stationary over time. By contrast, in the warped GP model, the inferred mean signal during the light step was smoother and tracked the faster oscillations much more closely (Fig. 2e). More importantly, in contrast to the interpolated signal derived by a classical pipeline, the warped GP infers a high level of uncertainty during periods of rapid oscillation which are at, or close to, the sampling limit of the recording.

### Using GP Models For Statistical Inference

The key benefit of the GP models is that they provide an explicit estimate of the uncertainty about the latent activity function which can be used to perform well calibrated statistical inference, e.g. for inferring which periods of neural activity differed between two conditions. This is in contrast to classical approaches, where typical analysis follows multiple smoothing steps and often only the inter-trial variability is considered, providing a poorly calibrated estimate of uncertainty.

In our framework, we use a GP Equality Test to identify whether two signals are statistically distinct [13]. As an example, we consider the response of bipolar cells to the chirp stimulus as a function of the spatial extent of the light spot. This is known to modulate bipolar cell responses, with the difference being induced by lateral inhibition [6, 14–16]. We compared the calcium and glutamate signals of bipolar cells presented with chirp stimuli whose light spots differed in size (100*μm* and full field). We fitted a GP model with time warping to each of the sets of observations (Fig. 4b), performed MLE optimisation and then computed the difference between the estimated latent functions.

**Fig. 4:**
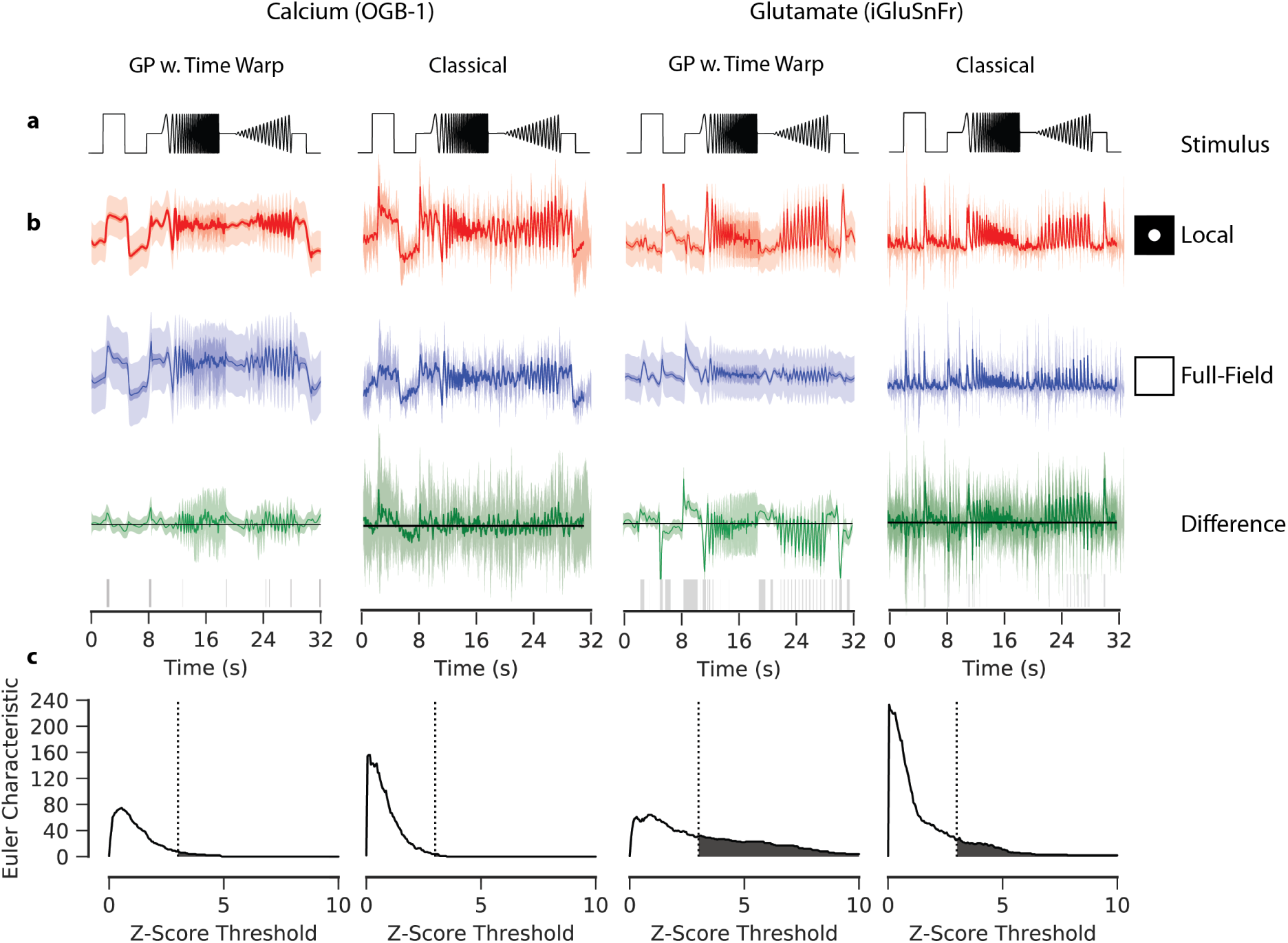
Gaussian Process Equality Testing. Data: ROI from a retinal bipolar cell filled with OGB-1 by sharp electrode injection (left), and a different ROI from a scan field with bipolar cell terminals in a retina expressing iGluSnFR (right); both recorded using spiral scan configurations. Model: GP w. Time Warp: RBF kernel, 300 inducing inputs, 20 iterations per fit, best of 5 fits per model. Classical: pipeline incorporating frame averaging and interpolation. (**a**) “Chirp” light stimulus. (**b**) Fits of the GP with time warping and classical pipeline to chirp-driven responses, for calcium and glutamate data. Models fitted to observations of the responses to local (100μm; top) and full field (middle) chirp stimulus. Circles indicate relative spatial extent of the light stimulus. Difference between the models for the two stimulus conditions shown at the bottom. Intervals for the response data show 3 standard deviations above and below the mean function. This variability corresponds to the standard deviation with and without additive noise for the GPs, and the inter-trial standard deviation for the classical pipeline. Only the standard deviation without additive noise is shown for the difference of the GPs. Domains where zero-vector not included within this interval are highlighted with grey ticks, corresponding to regions where the difference between the two signals is greater than expected by chance. (**c**) Frequency of discrete domains where zero-vector is not included in the credible intervals (also known as the Euler Characteristic; EC) as a function of the number of standard deviations above and below the respective mean functions. A high EC indicates a large degree of statistical separation between the two signals, and typically declines as the threshold increases.

In addition, in a practical context one wants to know not merely whether two functions differ, but also when. In the case of light driven neural activity, this corresponds to the periods of activity where the stimulus drives greater differences in the response than would be expected by chance. This we specified to be the discrete regions where the difference between the two latent activity functions was greater than three standard deviations, and were therefore statistically significant, following the approach used in [17]. We found that regions where the two latent activation functions were significantly different occurred during parts of the light step and in both oscillatory sequences.

The number of disconnected regions where the difference is greater than could be expected by chance is called the Euler characteristic, and provides a measure of the strength of the difference between two signals (Fig. 4c). As expected, we found the responses of the two bipolar cells to show clear evidence for differences between the two stimulation conditions.

For comparison, we computed a comparable test using the classical analysis pipeline, using inter-trial standard deviation as an estimate of the uncertainty associated with the mean signal. Here, the difference in the estimate of the uncertainty for the classical and GP approaches becomes critical: the classical approach does not distinguish between observational and stimulus driven noise. There are three reasons to consider such an approach inferior to our GP framework. Firstly, since there are usually very few trials the estimate of the variance at each time point is computed from an extremely small sample. Secondly, the estimate does not account for the statistical effect of the smoothing procedure which precedes it. Finally, the method does not consider autocorrelation - for the Euler Characteristic in particular this concern is critical, as the number of discrete regions where two signals differ is necessarily contingent on their autocorrelation characteristics. In contrast to the GP approach, the regions where the signals differ more than chance do not correspond to interpretable stimulus components, suggesting that they are merely a consequence of observation noise.

### Evaluating Hierarchical Clustering With GP Models

We next show how GP Equality Tests can be used to provide a principled criterion for choosing the number of clusters in a hierarchical clustering of light responses. For example, in a single imaging plane, one may wish to know whether the observed responses originate form distinct functional groups, perhaps due to the presence of multiple cells or cell types within the recording plane, or multiple neurites of the same cell acting independently (“multiplexing”, [18]).

We tested this by using GPs fitted to the activity of ROIs in GCaMP6f labelled bipolar cell axon terminals in a PCP2 mouse line (data previously published in [6]). We used the mean functions to compute a hierarchical clustering of the data (Fig. 5a). The model for each edge in the hierarchy was then fitted to the pooled observations of all the signals within the corresponding cluster. The split at each node was evaluated using a GP Equality Test (Fig. 5e,f,g) using again the Euler characteristic as an indicator of evidence for a significant difference.

**Fig. 5:**
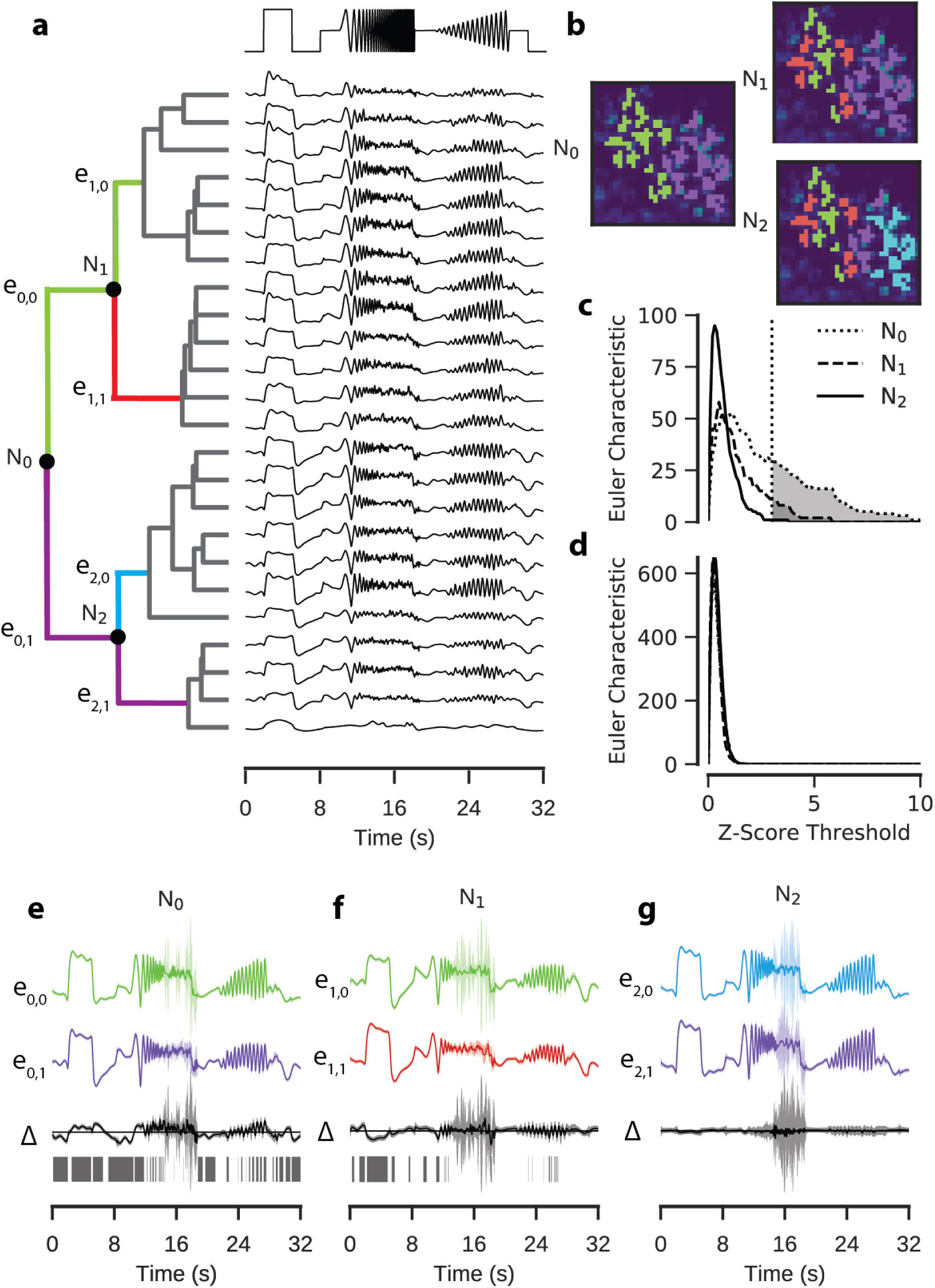
Hierarchical Clustering Of Retinal Bipolar Cell Terminals. Data: ROIs from a retinal bipolar cell filled with OGB-1 by sharp electrode injection recorded using a linear scan configuration. Model: RBF kernel (time), 300 inducing inputs, 20 iterations per fit, best of 3 fits per model. One model fitted for each ROI. (**a**) Mean functions of GP models fitted to calcium activity in a single recording field. Dendrogram computed using the Ward’s hierarchical clustering algorithm (left). Nodes where the equality test were performed are labelled N. Colours on the dendrogram correspond to putative clusters. (**b**) ROI masks overlaid on mean field activity, coloured with respect to the putative cluster. Each overlay is coloured according to the clustering at the corresponding nodes in (a). (**c**) Euler Characteristic for each node with respect to the z-score threshold. (**d**) Euler Characteristic for each node, using estimates of mean and variance from classical pipeline. (**e, f,g**) GP Equality tests performed at each of the labelled nodes. GPs correspond to the models fitted to each putative cluster (top, middle) and the difference between the two models (bottom). Intervals correspond to 3 standard deviations above and below mean function. Domains where zero-vector not included within interval highlighted with grey ticks.

Interestingly, the first split (N0) separates ROIs nicely into ROIs belonging to either of two bipolar cells in the imaging field, with strong quantitative backup for the split. Successive nodes down the hierarchy exhibited lower statistical evidence for the proposed clustering (Fig. 5c). Nevertheless, one further split of ROIs (N1) was accepted according to our statistical criterion, splitting ROIs of the left bipolar cell into two groups, indicating potential spatial variation within the terminals of a single bipolar cell. Subsequent separations were rejected.

We again compared the utility of using the GP models to a classical approach based on inter-repetition s.d. (Fig. 5d). As we had observed in our previous application of this classical equality test, observation noise obscured the stimulus-driven differences between the putative clusters. Consequently the test derived very similar estimates for the difference between the clusters, providing a poor measure of cluster quality.

### Incorporating Stimulus Effects Into GP Model Inference

We next extend our GP framework to study the effect of multiple stimulus parameters and their interactions on the latent neural activity in an ANOVA-like framework. In contrast to classical ANOVA, the interaction effects can have non-linear structure [19], and it is possible to compute not merely the strength of particular effects but also an inference of the response of a ROI over time as a stimulus feature varies. Unlike the previous two stages, there is no straightforward way to extend the classical analysis pipeline to deal with these more complex scenarios.

To demonstrate the usefulness of this extension, we fitted a GP model to a single ROI responding to a light stimulus where light intensity was modulated as a sine wave of different frequency and contrast (Fig. 6b). For the experiments, we selected a set of 90 stimulus parameters using blue noise sampling, such that parameters were uniformly selected from the parameter space and excluded if they were below certain thresholds for frequency (*<* 1*Hz*) or contrast (*<* 10%) or too close to an already existing stimulus parameter.

**Fig. 6:**
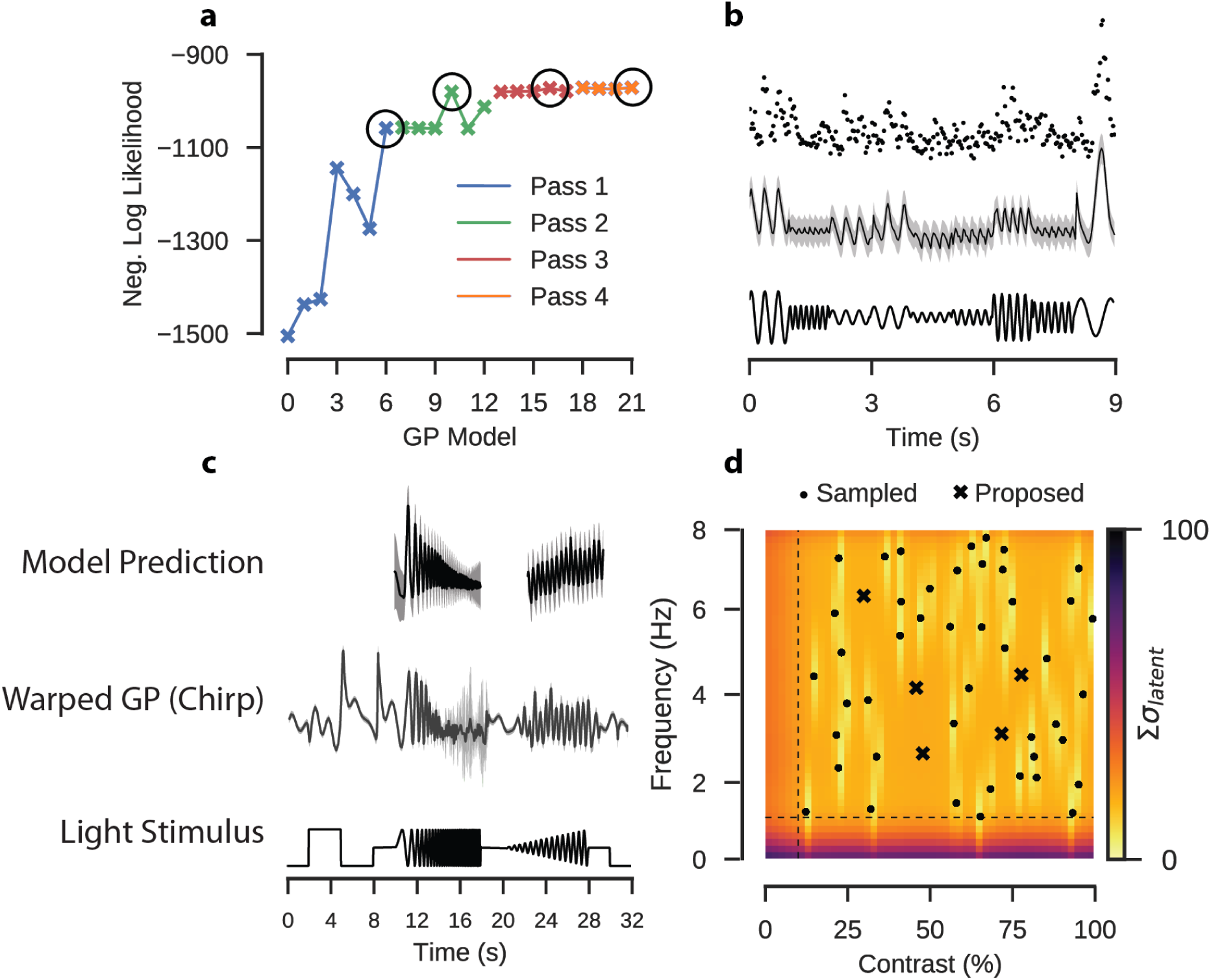
Extended Gaussian Process Model Incorporating Stimulus Parameters. Data: ROI from a scan field with bipolar cell terminals in a retina expressing iGluSnFR, recorded using a spiral scan configuration. Model: Product of RBF kernel (time) with composite RBF kernels (frequency and contrast), 500 inducing inputs, 25 iterations per fit, best of 3 fits per model. (**a**) Negative log likelihood for each model tested during model selection. (**b**) Observed activity of one ROI filled with OGB-1 (top); GP model selected by model selection procedure, conditioned on the observations of the ROI. Sine stimulus activity. Data corresponds to one 9 window of the stimulus. (**c**) GP fitted to observed chirp responses for the same ROI (middle). Prediction of the activity by the model on the sine stimulus data (top). (**d**) Locations in frequency-contrast parameter space selected for the stimulation. Colour map corresponds to the sum of the variance of the latent function for the fitted GP model evaluated under each parameter combination. Crosses correspond to peaks in the uncertainty where the stimulus should next be evaluated. Dashed line indicates the limits of the space from which the parameters were sampled.

As in a classical ANOVA, there are many possible ways in which the effects of these stimulus parameters can be incorporated into the model. In this case, stimulus features were encoded in the covariance kernel (see Methods), either as additive kernels for independent effects of phase, contrast or frequency, or through multiplicative kernels for interactions between the features. The cost of adding more kernels with a fixed observation set is that the uncertainty associated with each parameter increases as the number of parameters to be learned grows. To compensate for this, we performed kernel selection through a two stage iterative process. The first stage identified the kernel which, when included, most strongly improved model performance, as measured by the log marginal likelihood. Once there were two or more parameters, each new kernel had to contribute a greater improvement to the model performance than could be expected by chance, as established by a likelihood ratio test (see Methods). If a kernel was accepted it was retained in the model in the consecutive iterations.

We fitted a GP model to the glutamate signal of a single ROI in the IPL in response to the sine stimulus using this procedure (Fig. 6a,b). After four iterations the improvement in model performance was less than the required ratio and the process ceased. The kernels which were accepted included an interaction kernel between all three parameters, a phase-contrast kernel (Λ = 156.73, *p <* 0.001), and a phase-frequency kernel (Λ = 15.87, *p <* 0.001). In the final iteration an additive phase kernel was rejected (Λ = 2.36, *p* = 0.123). This suggests that there is a high level of interaction between the stimulus parameters.

We then used the model to predict neural activity for unseen parameter combinations and quantified how uncertain our predictions about the activity in response to these were [20–22]. Intuitively, the model should have the least uncertainty about stimulus parameters which had been observed. Uncertainty then should increase as a function of the distance from the observed parameters. We quantified uncertainty by computing the expected response of the ROI and taking the sum of the latent variance (Fig. 6d).

For the studied cell, calcium recordings to stimulation with the chirp stimulus were also available (Fig. 6c), and we compared the model fitted directly to the chirp response data to predictions from the model fitted to the sine data. There were some qualitative similarities between the two models, such as the overall amplitude of the signal, and the decrease in signal amplitude as the frequency of the stimulus increased. The prediction that the signal amplitude would slightly increase with contrast was not reflected in the chirp data, where the relationship was more ambiguous. The quality of prediction of the activity at high frequencies was difficult to evaluate, as there is a high-level of uncertainty about the mean signal at those frequencies. One factor to consider with regards to this direct comparison is that the differences in the chirp responses may be due to temporal dependencies over time.

### Active Bayesian Experimentation

A critical advantage of our framework is that we can use it for Bayesian experimental design. This is useful, as in 2P imaging experiments time is usually severely limited. For example, isolated mouse retinal tissue becomes unresponsive to light stimulation in a matter of hours, and single recording fields often bleach within half an hour of recording. To efficiently explore the space of possible stimulus features under severe time constraints is thus a critical problem, which GP models can be used to address [20, 21].

To show how this works, we performed an experiment using GP models to guide parameter selection. In retinal tissue expressing iGluSnFR we selected a single ROI, likely representing a single bipolar cell axon terminal. We used two control stimuli to evaluate the parameter selection: a local chirp stimulus playing over three trials, to which we fitted a warped GP, and a sinusoidal stimulus with 90 parameters uniformly sampled from the parameter space (Fig. 7a), to which we fitted a GP with the kernels derived in the previous likelihood ratio procedure. We performed three rounds of active parameter selection, starting with 30 uniformly sampled parameters in the first iteration, fitting the GP and using parameters selected by identifying 30 peaks in the uncertainty map in the subsequent two iterations. We then used the models from each iteration to predict how the ROI would respond during the oscillatory components of the chirp stimulus (Fig. 7b).

**Fig. 7:**
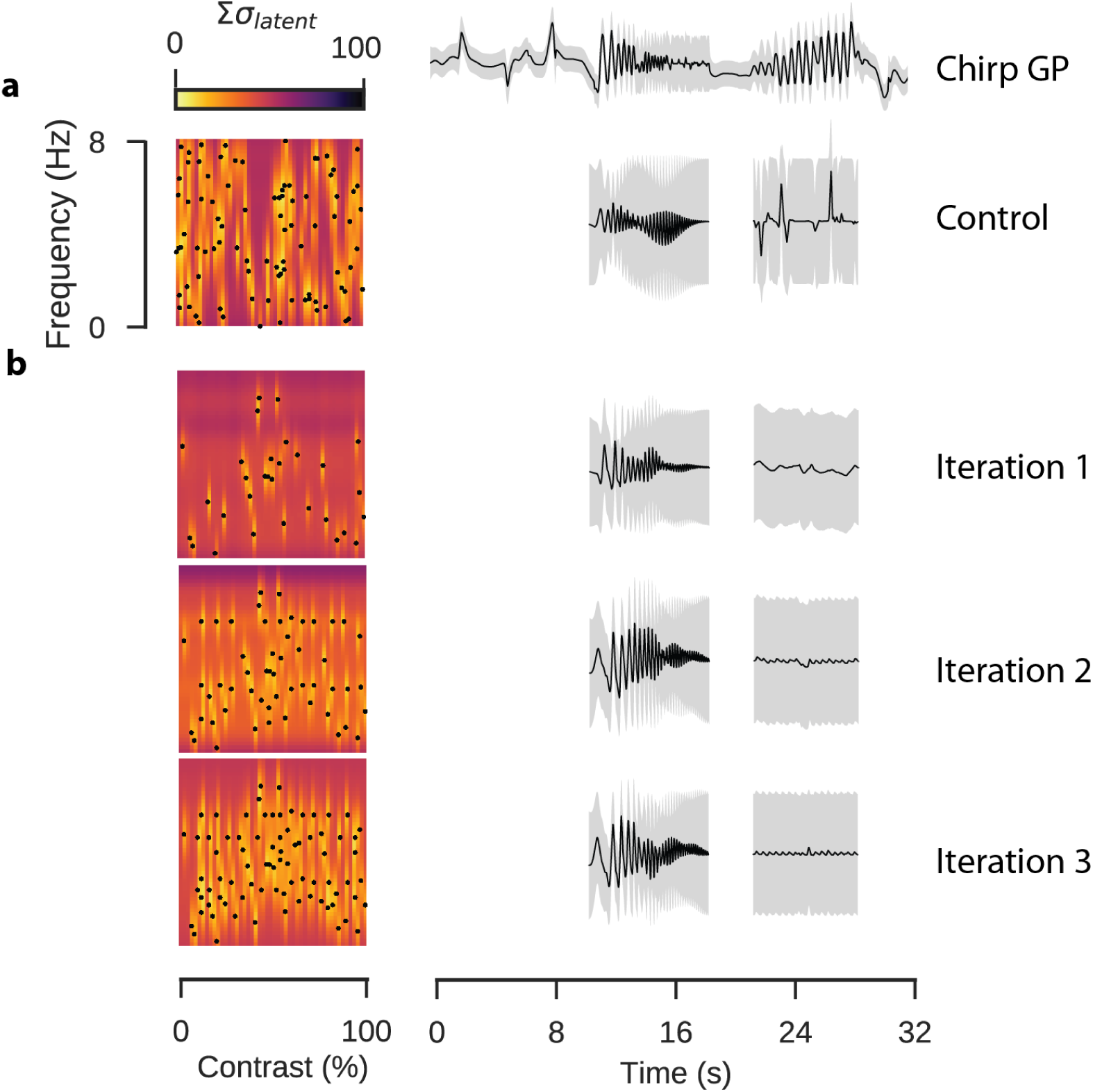
Active Parameter Selection With Gaussian Processes. Data: ROI from a scan field with bipolar cell terminals in a retina expressing iGluSnFR, recorded using a spiral scan configuration. Model: Product of RBF kernel (time) with composite RBF kernels (frequency and contrast), 500 inducing inputs, 25 iterations per fit, best of 4 fits per model. (**a**) Control stimulus consisting of 90 parameter sets of frequency and contrast. Uncertainty in each region computed as the sum of the latent uncertainty for a GP estimated under all parameter configurations. The chirp response for this ROI is shown above. The completed GP model for the sine response is estimated over the full dataset, the model inference for the sinusoidal chirp components is shown. (**b**) Active parameter selection using GP with corresponding inferences for chirp stimulus. The algorithm first separated ON and OFF responses into separate clusters. The ON cluster was then further divided into sustained and transient responses. The sustained ON responses were finally separated into orientation selective and non-orientation selective clusters. The strength of the evidence for a split of ON from OFF responses was greater than the subsequent partitions. The evidence for the separation of transient and sustained, or OS and non-OS, was roughly equivalent, suggesting a comparable level of evidence for each separation of the data into these clusters (Fig. 8f).

Parameters selected using the active approach were more broadly distributed across the parameter space, although we noted that the peak finding algorithm was biased away from the edges. In the purely random design procedure parameters often clustered and there were large empty regions, resulting in high uncertainty in these regions. Neither the random nor the active parameter procedure inferred a good prediction of the contrast-varying chirp component, which in the case of the active parameter inference was likely due to the bias away from the periphery of the parameter space, resulting in very few samples in the proximity of the 8Hz parametric edge. At lower frequencies the experimental design algorithm seemed better able to capture qualitative aspects of the chirp response, such as the decrease in response amplitude as the frequency increased, though again the lack of samples at the very highest frequencies resulted in a high level of uncertainty.

### Combining Model Components

We finally constructed a model which combined stimulus effect modelling and hierarchical clustering into a single framework. We fitted the model to calcium recordings of RGC activity in response to a bright bar moving in different directions on a dark background. RGCs show different response polarities and a large range of response kinetics to this stimulus [10] and some modulate the response amplitude as a function of stimulus direction. The model incorporated the stimulus features of time and direction as additive effects, alongside with a time-direction interaction effect (Fig. 8a,b). The data were then sorted using hierarchical clustering (Fig. 8c,d; Supplementary Fig. S3) and for the purpose of demonstration the first three nodes of the hierarchy were tested using GP Equality Tests (Fig. 8e).

**Fig. 8:**
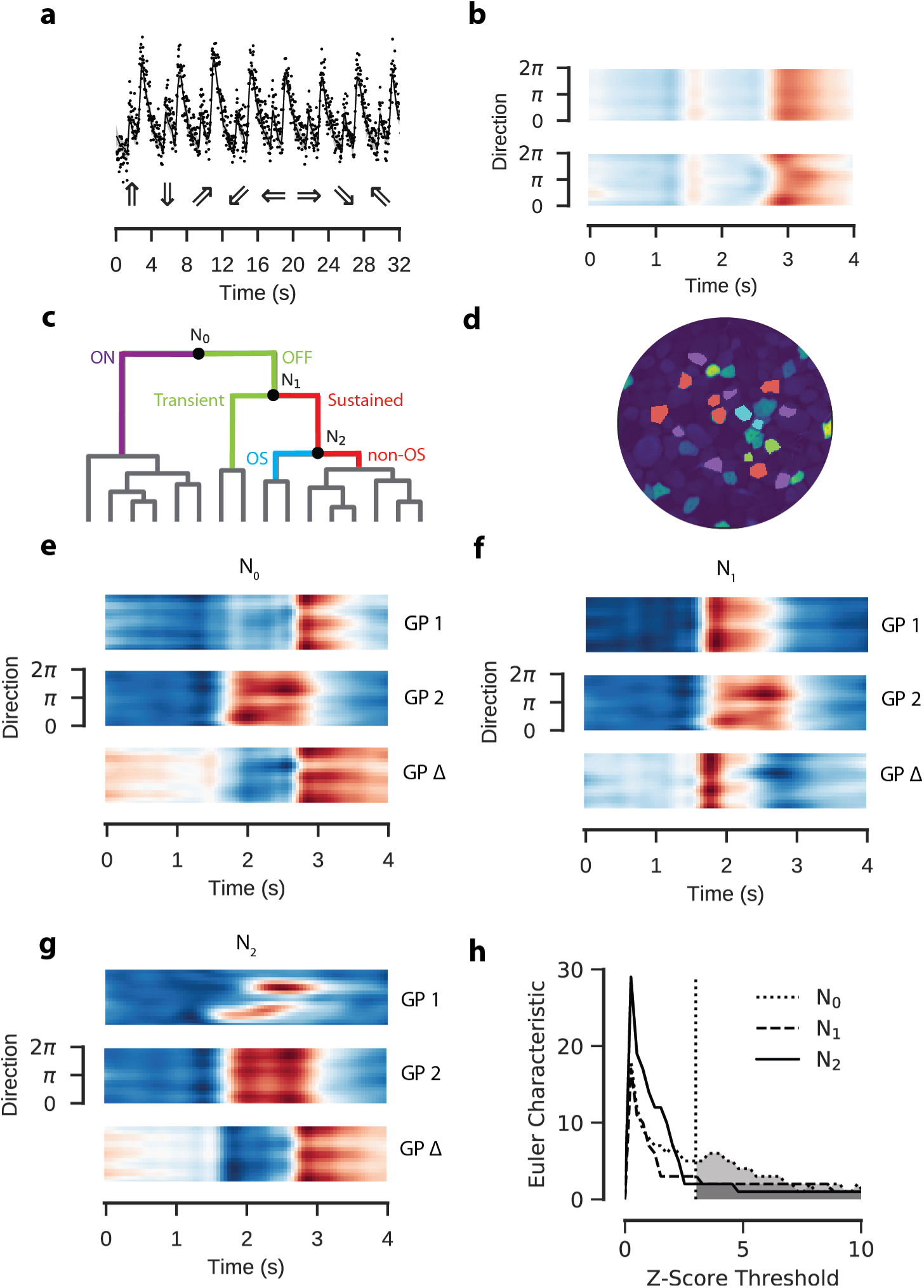
Gaussian Process Model Of Retinal Ganglion Cell Responses To A Moving Bar Stimulus. Data: ROIs from a field of RGC somata labelled with OGB-1 using electroporation, recorded using a spiral scan configuration. Model: Product of RBF kernel (time) with composite RBF kernel (orientation), 300 inducing inputs, 25 iterations per fit, best of 3 fits per model. (**a**) Observed activity of one ROI representing an RGC soma labelled with OGB-1 (top). GP model superimposed. Intervals correspond to the variance of the latent function, 3 standard deviations above and below the mean. Below are the moving bar directions for each trial. (**b**) GP model fitted to data in (a) without (top) and with (bottom) interaction kernel. Both models include additive effects for orientation and time. Coloured according to response amplitude (red: high, blue: low). (**c**) Hierarchical clustering of fitted models. Colours on dendrogram correspond to colours on ROI mask. Posterior means for each ROI in each cluster are shown in Supplementary Fig. S2. (**d**) ROI mask with cluster colours for k = 4 corresponding to clustering in (c). A baseline s.d. was computed from the 500ms of activity preceding the light step. ROIs where the amplitude of the step response was less than 2 s.d. greater or lower than the mean signal were excluded. (**e,f,g**) GP Equality Tests for each node. Top and middle correspond to the two putative clusters, bottom is the mean difference between them. (**h**) Euler Characteristic as a function of the number of standard deviations above and below the respective mean functions, for each node.

## Discussion

Here we presented a data analysis pipeline for 2P imaging data based on Gaussian Processes. The advantage of this framework is that uncertainty about the underlying latent neural activity can be propagated through the analysis pipeline, so statistical inference can be performed in a principled way. We applied our pipeline to recordings of mouse retinal bipolar and ganglion cell activity driven by light stimuli, showing how: to determine whether and when two response functions are statistically distinct; to evaluate the strength of the evidence for a partition of data into functional clusters; to determine the relevant stimulus effects to incorporate into a model of neural responses; and to guide the choice of stimulus parameters for iterations of a closed loop adaptive experiment.

### Estimation of Uncertainty

Accurately characterising the variability of neural responses is essential for understanding neural coding. Noise manifests itself throughout sensory systems and presents a fundamental problem for information processing [2]. While imaging *ex-vivo* retinal tissue does not present some of the challenges as *in vivo* cortical recordings (where movement is a significant source of variability), two-photon imaging in *ex-vivo* tissue is still subject to many sources of variance, due to fluctuations in biosensor excitation and photon detection, among other factors. Computational processing can introduce further variability, due for example to the discretisation of the measured signal. This is rarely acknowledged, perhaps due to the convenience of standard approaches.

### Computational Limitations of the Proposed Models

Classical GP models can be computationally costly due to the need to compute the inverse of the kernel matrix involving all training data [4]. To make practical use of GPs for modelling large 2P recordings, we capitalized on recent advances in sparse approximations for GPs that work with a limited number of inducing points [23] and demonstrated their applicability for a real world task. In addition, we only performed point-estimates for hyper-parameters instead of fully Bayesian inference and pre-determined kernels before statistical evaluation. This was important for two reasons: firstly, a processing pipeline should not be excessively computationally costly, so as to make them impractical for general use with larger datasets; and secondly, the application of these models in closed-loop imaging experiments was only possible if one complete iteration of the process (data acquisition; pre-processing; prediction; parameter selection) could be completed in a few minutes. In principle, our approach could be extended to a fully Bayesian framework with hyperpriors on the model parameters, although this introduces additional difficulties for sparse approximation and still entails a greater computational burden [24]. While our work solely addressed Gaussian distributed data, the models can be readily extended to point processes as well. There, sparse approximation techniques overcome the computational intractability of the model, and allow inference on relatively large datasets [25].

### Active Experimental Design of 2P Experiments

Although we demonstrated the potential for using GP models during 2P experiments, there were several limitations to our approach. We were able to reduce the time per iteration of our active experiments to less than five minutes, addressing a key practical concern. However, it emerged during the experiment that the peak-finding algorithm was biased away from the periphery of the stimulus space, which made the chirp stimulus unsuitable as our “ground truth” for model evaluation.

The parameter batch size may also have been too small for each iteration. Batch size is a critical consideration in active Bayesian experimentation. Where the cost per iteration is low, single parameters can be selected for each iteration, for which the objective function can be relatively easily defined and evaluated. In one recent publication, Charles and co-workers [22] used GPs to model the effect of inter-trial variability in monkey V1 neurons, using sequential parameter selection to optimise a coloured light stimulus. For experiments where iterations are prohibitively expensive, new parameters can be selected in batches, although this requires interactions between parameters to be taken into account, which can be computationally expensive to evaluate. In such cases, approximate methods provide an attractive method for reducing computational overheads (e.g. [21]). Batch parameter selection algorithms which account for, or approximate, parameter interactions would likely overcome simple peak finding methods.

### Conclusion

Historical obstacles to the use of Bayesian methods such as the difficulty of their implementation and their computational cost have been reduced. Much research over the past decade has focused on the problem of minimising the computational complexity of the algorithms through sparse approximation methods and efficient parameter estimation (such as [23]). New libraries for popular coding languages - such as GPy [26], PyMC3 [27], and pySTAN [28] for Python 3 - have lowered the barrier to entry. Likewise, we provide a collection of notebooks with this paper to allow straightforward application of our framework. Taken together, our approach exploits the flexibility and extensibility of Gaussian process models to improve on classical approaches and address important analytical tasks. We feel that it will be particularly useful for disentangling the dynamics of neural circuits in the early visual system under complex, multivariate experimental conditions.

## Methods

### Animals And Tissue Preparation

All animal procedures were performed according to the laws governing animal experimentation issued by the German Government. For single-cell-injection experiments, we used one adult mouse cross-bred between transgenic line Cg-Tg(Pcp2-cre)3555Jdhu/J (Tg3555, JAX 010536) and the Cre-dependent red fluorescence reporter line Gt(ROSA)26Sor^tm9(CAG-tdTomato)Hze^ (Ai9^tdTomato^, JAX 007905). For glutamate-imaging, we used one adult C57BL/6 mouse. Owing to the exploratory nature of our study, we did not use blinding and did not perform a power analysis to predetermine sample size.

Animals were housed under a standard 12h day-night cycle. For recordings, animals were dark-adapted for ≤ 1h, then anaesthetised with isoflurane (Baxter) and killed by cervical dislocation. The eyes were removed and hemisected in carboxygenated (95% O2, 5% CO_2_) artificial cerebral spinal fluid (ACSF) solution containing (in mM): 125 NaCl, 2.5 KCl, 2 CaCl_2_, 1 MgCl_2_, 1.25 NaH_2_PO_4_, 26 NaHCO_3_, 20 glucose, and 0.5 L-glutamine (pH 7.4). Then, the tissue was moved to the recording chamber of the microscope, where it was continuously perfused with carboxygenated ACSF at ∼ 37 ^°^C. The ACSF contained ∼ 0.1 *μ*M sulforhodamine-101 (SR101, Invitrogen) to reveal blood vessels and any damaged cells in the red fluorescence channel. All procedures were carried out under very dim red (*>*650nm) light.

### Single Cell Injection

Sharp electrodes were pulled on a P-1000 micropipette puller (Sutter Instruments) with resistances between 70 - 100MΩ. Afterwards, the tip (∼ 500*μ*m) of each electrode was bent on a custom-made microforge. Single bipolar cell somata in the inner nuclear layer were filled with the fluorescent calcium indicator Oregon-Green BAPTA-1 (OGB-1) by using the pulse function (500ms) of the MultiClamp 700B software (Molecular Devices). OGB-1 (hexapotassium salt; Life Technologies) was prepared as 15mM in distilled water. Immediately after filling, the electrode was carefully retracted. Imaging started after about 30 minutes after the injection to allow cells to recover and the dye to diffuse within the cell. At the end of the recording, a stack of images was captured for the cellular morphology, which was then traced semi-automatically using the Simple Neurite Tracer plugin implemented in Fiji [29].

### Virus Injection

For virus injections, we used adult wild-type mice (C57BL/6). Animals were anesthetized with 10% ketamine (Bela-Pharm GmbH & Co. KG) and 2% xylazine (Rompun, Bayer Vital GmbH) in 0.9% NaCl (Fresenius). A volume of 1*μ*l of the viral construct (AAV2.hSyn.iGluSnFR.WPRE.SV40, Penn Vector Core) was injected into the vitreous humour of both eyes via a Hamilton injection system (syringe: 7634-01, needles: 207434, point style 3, length 51mm, Hamilton Messtechnik GmbH) mounted on a micromanipulator (World Precision Instruments). Imaging experiments were performed 3 weeks after virus injection.

### Two-Photon Imaging

We used a MOM-type 2P microscope (designed by W. Denk, now MPI Martinsried; purchased from Sutter Instruments/Science Products). The design and procedures have been described previously [6, 10, 30]). In brief, the system was equipped with a mode-locked Ti:Sapphire laser (MaiTai-HP DeepSee, Newport Spectra-Physics), two fluorescence detection channels for OGB-1 or iGluSnFR (HQ 510/84, AHF/Chroma) and SR101/tdTomato (HQ 630/60, AHF), and a water immersion objective (W Plan-Apochromat 20x /1.0 DIC M27, Zeiss). The laser was tuned to 927nm for imaging OGB-1, iGluSnFR or SR101. For image acquisition, we used custom-made software (ScanM by M. Müller, MPI Martinsried, and T. Euler) running under IGOR Pro 6.3 for Windows (Wavemetrics), taking time lapsed 32 x 32 pixel image scans (at 15.625Hz) or 16-line “spiral” scans (at 31.25Hz). For documenting morphology, 512 x 512 pixel images were acquired with step size of 0.5*μ*m along the Z axis.

### Fast Spiral Scan Imaging

To resolve transient changes in calcium concentration or glutamate release (i.e. with decay times of ∼100ms), scan rates of around 20Hz or more are wanted. Many scanning 2P microscopes use conventional (non-resonant) galvanometric scanners and are limited by the inertia of the scan mirrors, which introduce positional errors at high scan rates. This is especially critical for typical linear (image) scans, with their abrupt changes in direction when jumping between scan lines. For constant spatial resolution, faster scan rates are often realised by decreasing the scan area. However, it is possible to increase the spatio-temporal resolution by using non-linear “spiral scan” configurations (Fig. 1c). These overcome the key mechanical limitation of linear scans, that they incorporate sharp turns, rather than following smoother trajectories. Unlike linear scans, which are composed of single linear trajectories repeated along an axis at regular intervals (Fig. 1d), spiral scan configurations consist of radial trajectories moving away from a central point at a constant speed and rotation and permit rapid movement of the scan mirrors.

A regular radial grid can be constructed by generating a single spiral trajectory and successively rotating it around a central point. We used an Archimedean spiral is used to generate each trajectory (*r* = Θ^1/*a*^), where the radial distance *r* from the central point is a function of the angle Θ and a tightness parameter *a* which determines the rate of rotation around the centre. With a grid composed of 16 such curves we can resolve, for instance, axon terminals of retinal bipolar cells at twice the spatial and twice the temporal resolution of linear recordings. One can see the advantages of such scan configurations by showing how frequently the scan trajectory intersects with ROIs in a single frame (Fig. 1e,f). The times at which labelled structures are observed by these trajectories are both more frequent and more irregularly distributed in time than a typical linear scan (Fig. 1g,h), providing a superior temporal resolution.

### Light Stimulation

For light stimulation, a modified LightCrafter (DLPLCR4500, Texas instruments; modification by EKB Technology) was focused through the objective lens of the microscope. Instead of standard RGB light-emitting diodes (LEDs), it was fitted with a green (575nm) and a UV (390nm) LED for matching the spectral sensitivity of mouse M- and S-opsins [31]. To prevent the LEDs from interfering with the fluorescence detection, the light from the projector was band-pass-filtered (ET Dualband Exciter, 380-407/562-589, AHF) and the LEDs were synchronised with the microscope’s scan retrace. Stimulator intensity was calibrated to range from 0.5 * 10^3^ (“black” background image) to 20 * 10^3^ (“white” full field) photoisomerisations P*/s/cone [6]. The light stimulus was centred before every experiment, such that its centre corresponded to the centre of the recording field. In linear scans, the stimulus is displayed while the trajectory moves between consecutive lines; while for the spiral scans this occurs while the trajectory returns from the periphery to the centre.

Light stimuli were generated using the QDSpy light stimulation software, which is written in Python 3 [32]. The chirp stimulus ran for 4 repeats of 32s each, with the stimulus extent alternating between a 800*μm* and a 100*μm* light spot. The moving bar stimulus consisted of a 300*μm* rectangular bar moving at 1000*μm/s* for 4 seconds along 8 evenly space orientations, repeated three times for each orientation. The sine stimulus consisted of a 100*μm* light spot, and ran for 45 1s-trials, with contrast and frequency varying in each trial. The contrast and frequency parameters were chosen by blue-noise sampling 90 parameters from the parameter space, between 10% and 100% contrast and 1Hz to 8Hz frequency. Later closed-loop experiments used a sine stimulus with 90 parameter sets sampled uniformly from the parameter space, in addition to 3×30 parameters sets, of which the first were chosen from random uniform sampling and the latter two sets by active Bayesian inference.

## Data Analysis

Initial data analysis was performed in Igor Pro 6. Regions of Interest (ROIs) were defined manually using the SARFIA toolbox for Igor Pro [33]. In the iGluSnFR recordings, a custom-script generated a correlation map (see methods in [6]), which defined structures for the ROI drawing. To reduce the dimensionality of these observation sets, we computed the mean observation of the ROI activity on each scan line. The observations were synchronised to the light stimuli using time markers which were generated by the stimulation software and acquired during imaging.

Once the initial pre-processing was completed, the data was exported to HDF5 files [34], and all subsequent analysis was performed in Python 3.5. The Gaussian process models were developed in the GPy framework [26]. Feature encoding, input warping, equality tests, parameter selection and closed-loop parameter selection were computed using custom scripts, which will be provided as supplementary content to this document. Hierarchical clustering was performed using scripts from the Scipy library, using Euclidean distance, the ward algorithm and maxclust as the criteria. Adaptive parameter selection used a local peak finding algorithm from the Scikit-Image library.

### Mathematics Of Gaussian Process Regression

We model our observations as a set of random variables corrupted by additive Gaussian noise. A further index set locates these variables with respect to our predictors, such as time or stimulus frequency. Any finite subset of these observations are assumed to form a multivariate Gaussian distribution, drawn from a latent Gaussian process.

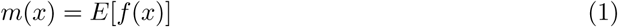

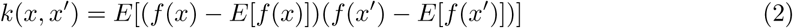

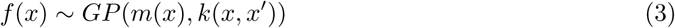

The Gaussian process is defined by its mean and covariance function. The covariance function describes the covariance between any two random variables in our observation set, and varies as a function of distance with respect to the predictors. It can be considered as a prior distribution over functions.

We used a radial basis function (RBF) kernel to model the covariance over the predictors. RBF kernels are members of the class of Matérn covariance kernels. The parameters of these covariance functions vary the smoothness *v*, length-scale *l*, signal variance *σ*_*signal*_ and noise variance *σ*_*noise*_. Several common basis functions correspond to Matrn functions with particular values for *v*, whose derivatives have closed forms which make them efficient to compute. Consequently we pre-specify *v.* The value of the parameters is learned as part of the optimisation.

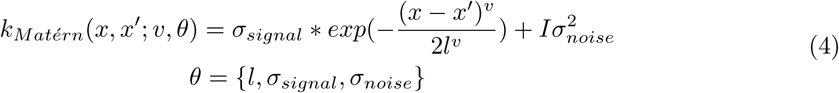

Gaussian processes define a joint distribution over the observation set and further test sets. As no two ROIs are actually observed at the same time, the joint distribution is used to infer estimates of the activity at a new, common set of test points, which can be used for statistical inference. By deriving the conditional distribution, we see that an intuitive explanation for the effect of the observations is that it subtracts uncertainty from the prior covariance.

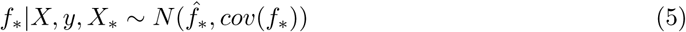

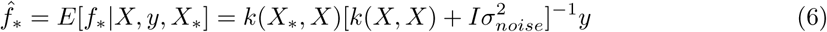

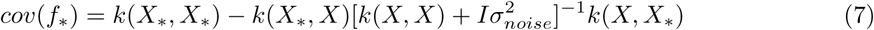

The log marginal likelihood is used as an objective function to define the quality of the model with respect to the observations. This can be derived by integration:

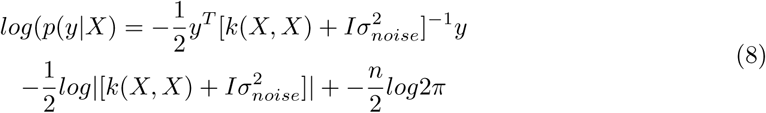

Evaluating the log marginal likelihood requires a matrix inversion which is cubic in computational complexity *𝒪*(*N* ^3^). To overcome this prohibitive cost, a sparse approximation *Q* of the covariance matrix *K* is used, whereby the covariance matrix is estimated over a subset of the observation set. The data in this subset are referred to as the inducing inputs.

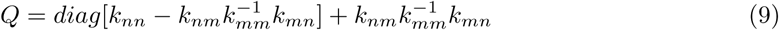

The choice of the subset of inducing inputs is learned as part of the optimisation. For this, the Kullback-Liebler divergence is used to define a distance measure between the exact Gaussian process and its approximation. We use the implementation in GPy, which follows from the algorithm of Titsias [23]. The parameters are optimised using the L-BFGS-B algorithm.

### Gaussian Process Equality Tests

The Gaussian Process Equality Test establishes whether two functions modelled by Gaussian processes are equal [35]. It operates by computing the difference between the two distributions and identifying whether the credible region encompasses the zero vector across the complete domain of the predictors. If the zero vector is outside of these intervals, we say the two functions are distinct with probability 1 - *a*. The probability is calculated using the mean *μ*^*^ and covariance *k*^*^ of the posterior of our two functions.

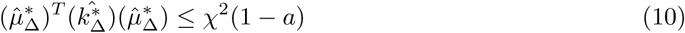

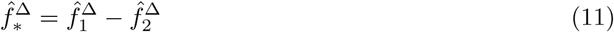

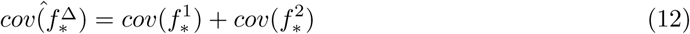

### Non-Stationary Autocorrelation

To address non-stationarity of the chirp response data, we computed the autocorrelation function for the chirp stimulus in 500ms windows spaced at 1/16s intervals (512 windows total). As we used RBF kernels for our regression, we fitted a Gaussian curve to each autocorrelation function and retained the inferred lengthscale *l*_*t*_ for each window. A further parameter *A* modulates the height of the function.

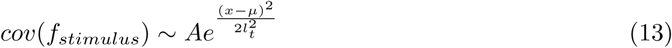

If the signal were stationary, we would observe that the lengthscale parameter was constant with respect to time. By using the cumulative distribution of the inverse of the lengthscale as the predictor, we could derive a warping function which transformed the predictors such that the stimulus autocorrelation was stationary. We assumed that the autocorrelation of the observed signal was approximately equal to that of the light stimulus input, and used the warping function to transform the observations. This transformation could be inverted to visualise the fitted gaussian process with respect to the original time base.

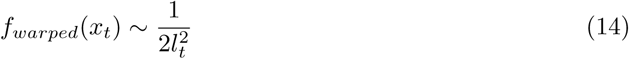

### Kernel Construction

Kernels can be combined in a number of ways [4], each expressing some belief about the effect of a parameter, most commonly by taking the sum or product of two kernels. Additive components represent effects of predictors which are independent of one another, while multiplicative kernels represent interactions between predictors.

For the chirp stimulus, where there is one predictor, a single kernel encoding the autocorrelation of the signal over time was used. For the moving bar and sine wave stimuli, additional kernels were included to model the effects of their respective parameters. For the moving bar the additional parameter is orientation, which was encoded as a 2D circular feature by converting the angle in polar coordinates to an xy position in Cartesian coordinates. The GP model for the moving bar responses included both additive effects for time and orientation, and a time-orientation interaction effect.

Circular features were also used to encode the phase of the sine stimulus, while frequency and contrast were encoded linearly. Likelihood ratio tests were used to select kernels from the full set of additive and multiplicative stimulus effects:

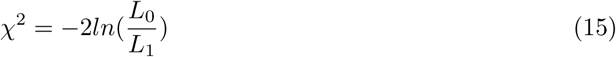

Where *L*_*N*_ is the likelihood of the fitted model, and the addition of the proposed parameter is rejected if the improvement in the likelihood is greater than chance with probability 1 - *a*. These tests were applied iteratively until a kernel was rejected. For the closed loop experimentation, we retained the model from the previous selection procedure with the randomly parameterised sine stimulus.

The data used throughout this paper and corresponding code used to compute the models will be provided as supplementary material alongside this paper.

## Supplementary Material

**Fig. S1:**
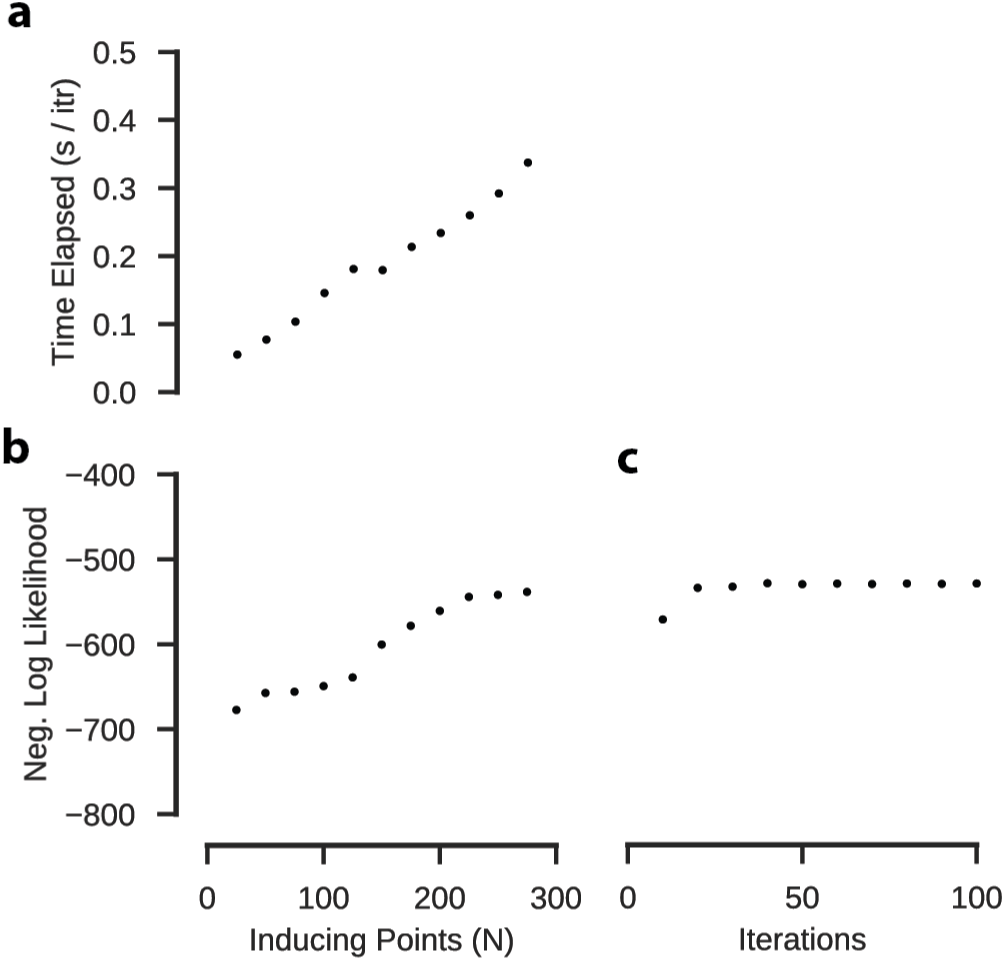
Model Performance Model: RBF kernel, best of 5 fits per model. (**a**) Mean time elapsed per iteration of the MLE. This scales approximately linearly with the number of points. (**b**) Out of sample estimate of negative log likelihood after 30 iterations. Estimates as a function of the number of inducing points. (**c**) Model performance of out of sample test points relative to the maximum number of iterations, evaluated for 300 inducing points.

**Fig. S2:**
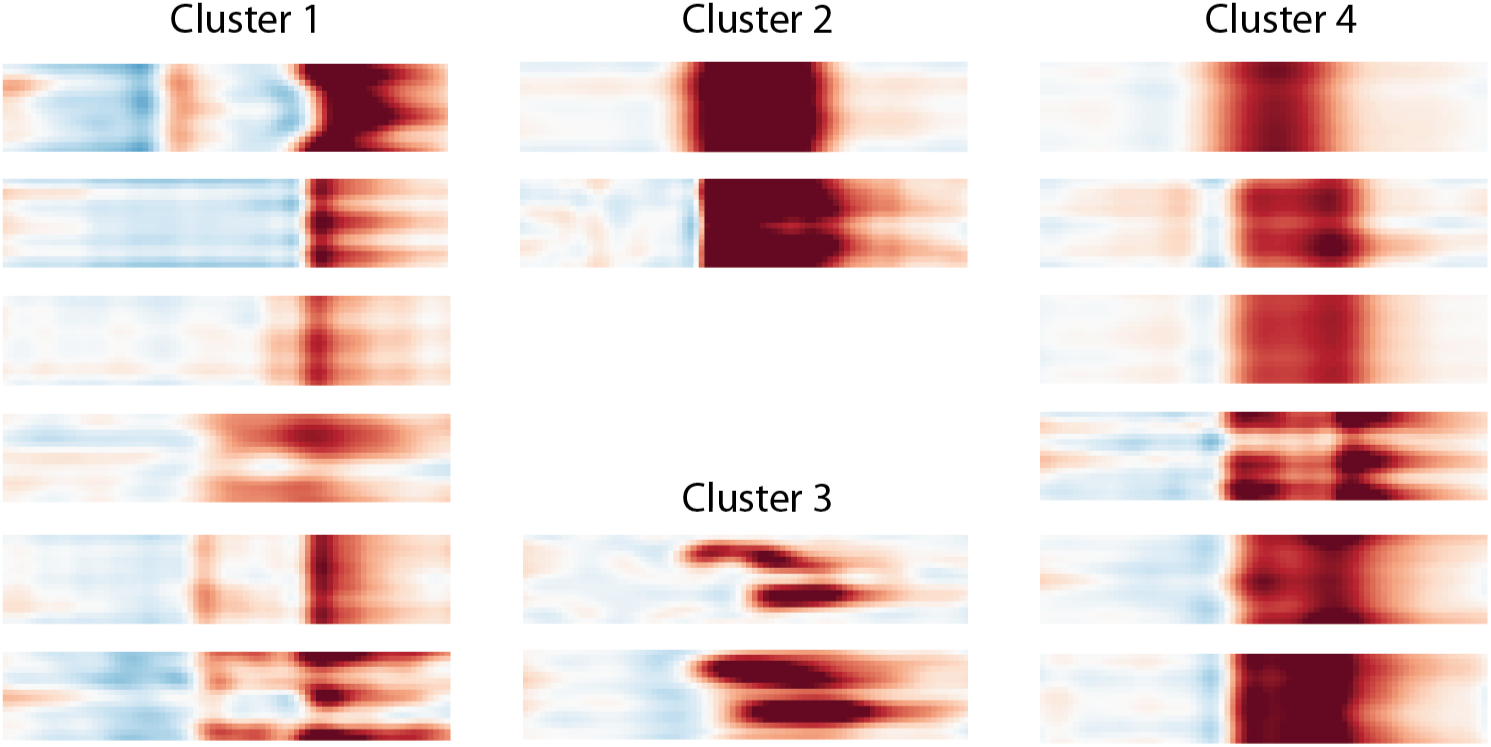
Direction Selectivity Clusters Model: Product of RBF kernel (time) with RBF kernel (orientation), 300 inducing inputs, 25 iterations per fit, best of 3 fits per model. Each heat map corresponds to the posterior mean of the fitted GP. Columns correspond to clusters.

### Oscillatory Kernels

| Kernel ID | Effects |
| --- | --- |
| 1 | Time |
| 2 | Freq |
| 3 | Cont |
| 4 | Time-Freq |
| 5 | Time-Cont |
| 6 | Freq-Cont |
| 7 | Time-Freq-Cont |

### Oscillatory Models

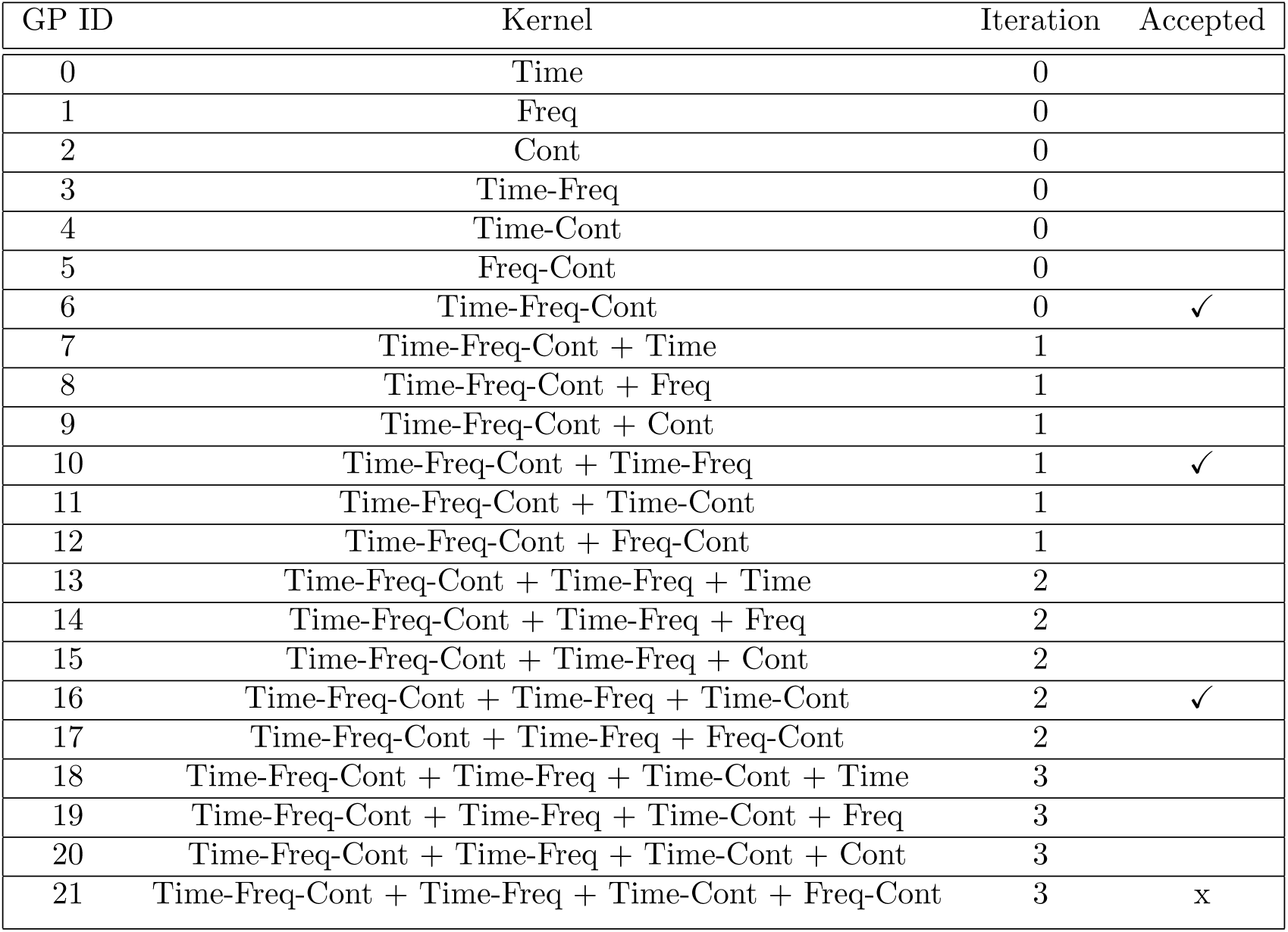

## Author Contributions

LER, TE and PB conceptualised the project, developed the methodology, and drafted the manuscript. LER, ZZ, and KF conducted the experimental investigation. LER performed formal analysis, implemented the software, validated the models, and visualised the results, with input from PB and TE. TE and PB provided funding, supervision, resources and project coordination. All authors contributed to the reviewing and editing of the manuscript.

## Acknowledgements

We thank Rob Smith and Roland Taylor for early inspiration; André Maia Chagas and Tom Baden for helpful discussion and feedback. We thank the authors of the GPy library, which provided a starting point for our model implementation.

